# Increasing Host Cellular Receptor—Angiotensin-Converting Enzyme 2 (ACE2) Expression by Coronavirus may Facilitate 2019-nCoV Infection

**DOI:** 10.1101/2020.02.24.963348

**Authors:** Pei-Hui Wang, Yun Cheng

**Affiliations:** Advanced Medical Research Institute, Shandong University, Jinan, Shandong 250012, China; School of Biomedical Sciences, The University of Hong Kong, Pokfulam, Hong Kong

**Author notes:** To whom correspondence should be addressed (P-H. W.).

## Abstract

The ongoing outbreak of a new coronavirus (2019-nCoV) causes an epidemic of acute respiratory syndrome in humans. 2019-nCoV rapidly spread to national regions and multiple other countries, thus, pose a serious threat to public health. Recent studies show that spike (S) proteins of 2019-nCoV and SARS-CoV may use the same host cell receptor called angiotensin-converting enzyme 2 (ACE2) for entering into host cells. The affinity between ACE2 and 2019-nCoV S is much higher than ACE2 binding to SARS-CoV S protein, explaining that why 2019-nCoV seems to be more readily transmitted from the human to human. Here, we reported that ACE2 can be significantly upregulated after infection of various viruses including SARS-CoV and MERS-CoV. Basing on findings here, we propose that coronavirus infection can positively induce its cellular entry receptor to accelerate their replication and spread, thus drugs targeting ACE2 expression may be prepared for the future emerging infectious diseases caused by this cluster of viruses.

## Introduction

The outbreak of a novel coronavirus (2019-nCoV) was reported in Wuhan, Hubei, China. According to the World Health Organization on February 24th 16, about 77,269 viral infected patients and at least 2596 deaths were confirmed globally. 2019-nCoV genome shows high similarity some bat coronaviruses as well as severe acute respiratory syndrome coronavirus (SARS-CoV) and Middle East respiratory syndrome coronavirus (MERS-CoV) which has caused two large-scale pandemics in the last two decades (Zhou et al., 2020). They all belong to betacoronavirus genus, which is a type of positive-stranded RNA viruses (Wu et al., 2020a; Zhou et al., 2020). Like SARS-CoV and MERS-CoV, 2019-nCoV can spread from the human to human and cause fever, severe respiratory illness, and a series of unidentified pneumonia disease (Chan et al., 2020; Huang et al., 2020; Li et al., 2020; Wang et al., 2020). Comparing with SARS-CoV and MERS, 2019-nCoV seems to be more readily transmitted from human to human, spreading to multiple countries and regions and leading to the WHO declaration of a Public Health Emergency of International Concern (Chan et al., 2020; Li et al., 2020; Wu et al., 2020b).

Coronaviruses utility the spike (S) proteins to select and enter target cells. SARS-CoV S protein and 2019-nCoV S protein engages angiotensin-converting enzyme 2 (ACE2) as their entry receptor (Li et al., 2003; Zhou et al., 2020; Wrapp et al., 2020). ACE2 is the cellular entry receptor of SARS-CoV and 2019-nCoV, which plays an indispensable role for SARS-CoV infection (Li et al., 2003; Zhou et al., 2020). ACE2 is a type I membrane protein expressed in the lung, heart, kidney, and intestine (Zhang et al., 2020). Enhanced expression of ACE2 accelerates SARS-CoV infection and spread, while silencing of ACE2 blocks its entry into cells (Li et al., 2003). Organs and tissues with high ACE2 abundance like the lung, kidney and small intestine are the infection targets of 2019-nCoV (Zhang et al., 2020). Taken together, these findings reveal that the expression level of ACE2 is extremely important for the successful infection of SARS-nCoV and 2019-nCoV. Here we analyzed the various microarray database of virus infected- or inflammatory stimulated-animals and cells, and clearly show that virus infection including SARS-CoV and MERS-CoV can obviously upregulate ACE2 expression, which suggests that 2019-nCoV may use similar strategies to augment its infection and spread.

## Materials and Methods

### Cell Culture and Transfection

HEK293 and HEK293T were cultured in Dulbecco’s modified Eagle’s medium (DMEM, Gibco, USA) with 10% heat-inactivated fetal bovine serum (FBS, Gibco, USA). poly (I:C) (Sigma P1530) was transfected into cells using Lipofectamine 2000 (Thermo Fisher, USA).

### Plasmids

pcDNA6B-TBK1-FLAG and pcDNA6B-TRIF-FLAG were constructed using standard molecular cloning methods as described previously (Wang et al., 2018).

### RT-qPCR

Total RNA was isolated by TRIzol® reagent (Invitrogen, USA). First-strand cDNA was synthesized with Transcriptor First Strand cDNA Synthesis Kit (Takara, Dalian, China). RT-quantitative PCR (RT-qPCR) reactions and analysis were performed using Roch LightCycler96. SYBR Green-based RT-qPCR were used according to the manuals by SYBR Premix Ex Taq kit (Takara Bio). Primers used in RT-qPCR reactions as follows. GAPDH-F(5’-3’) : GGAGCGAGATCCCTCCAAAAT; GAPDH-R(5’-3’) : GGCTGTTGTCATACTTCTCATGG. ACE2-F(5’-3’): GGAGTTGTGATGGGAGTGAT; ACE2-R(5’-3’)GATGGAGGCATAAGGATTTT.

### Virus Infection

VSV-GFP infection was described in our previous publications (Wang et al., 2018). Briefly, target cells were washed twice with 37°C DMEM medium, then virus that diluted in serum-free medium were added to the cells. Virus-medium complexes were incubated with cells in normal culture condition. 1-2 hrs later, virus-medium complexes were discarded and replaced with culture medium containing 10% FBS.

### Statistical Analysis

Three independent experiments are performed in all studies. Two-tailed unpaired Student’s t test by GraphPad Prism 7.0 was used to perform statistical analysis. P < 0.05 was considered as statistically significant in all experiments.

## Results

### Transfection of Poly (I:C) Induces ACE2 Expression

poly (I:C) is mimics of virus RNA. When delivered into HEK293 cell, human ACE2 mRNA expression level is significantly enhanced by RT-qPCR study (Figure 1). poly (I:C) mimics virus RNA and activate immune signaling pathways which lead to interferons (IFN) and inflammatory cytokine production, so next we explored whether RNA virus infection can induce ACE2 expression. As an RNA virus, VSV-eGFP can slightly induce ACE2 mRNA upregulation in HEK293 cells (Figure 2).

**Figure 1.**
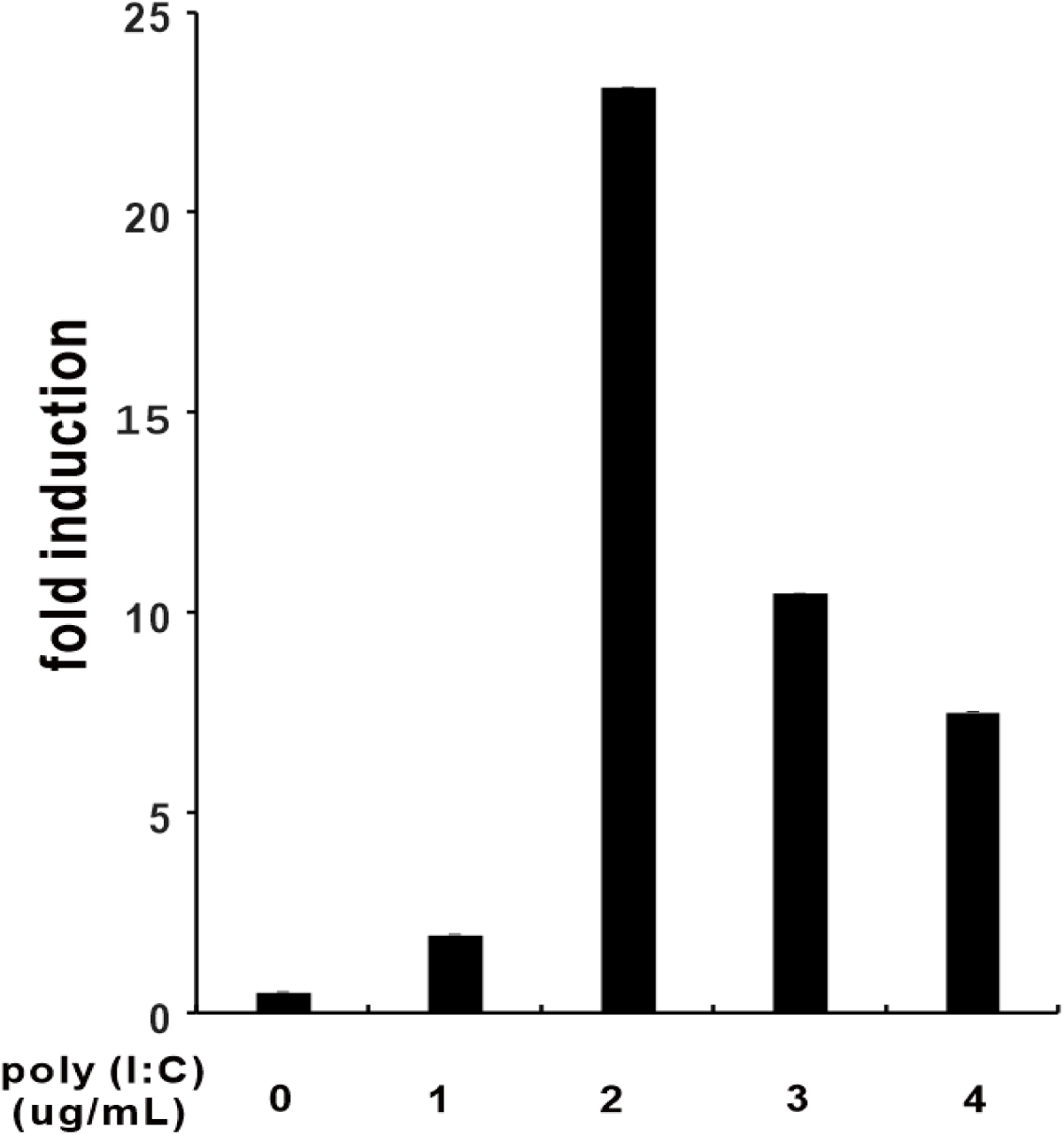
Transfection of poly (I:C) induces ACE2 expression. HEK293 cells were cultured in six-well plates and transfected with poly (I:C) as indicated above using Lipofectamine 2000. HEK293 cells were then collected for RNA isolation and RT-qPCR analysis at 9 hrs post-transfection. Results in each panel are representative of three independent experiments.

**Figure 2.**
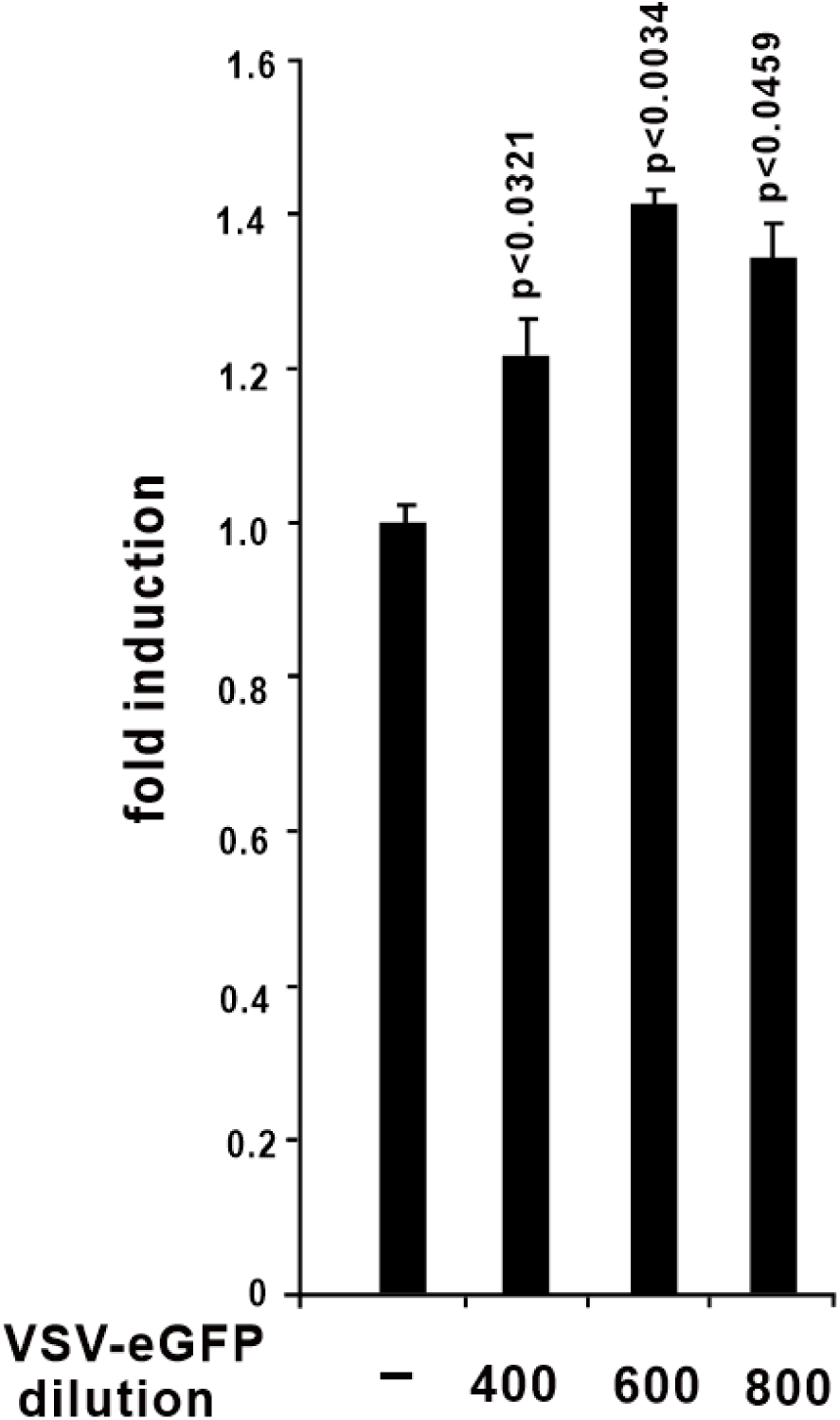
VSV-eGFP infection induces ACE2 expression. HEK293 cells were cultured in six-well plates and infected with VSV-eGFP as indicated above. 6 hrs later, HEK293 cells were collected for RNA isolation and RT-qPCR analysis. Results in each panel are representative of three independent experiments.

### Activation of RNA-sensing Pathway Induces ACE2 production

Because both poly (I:C) and RNA virus induce ACE2 expression, so next we evaluate whether RNA-sensing pathway have effect on ACE expression. Overexpression of TBK1, downstream of both TLR3 and RIG-I-like receptor signaling pathway, can strongly upregulate ACE2 mRNA level (Figure 3). TRIF, an adaptor protein of TLR3 pathway, also induce the expression of ACE2 in HEK293 cells (Figure 3).

**Figure 3.**
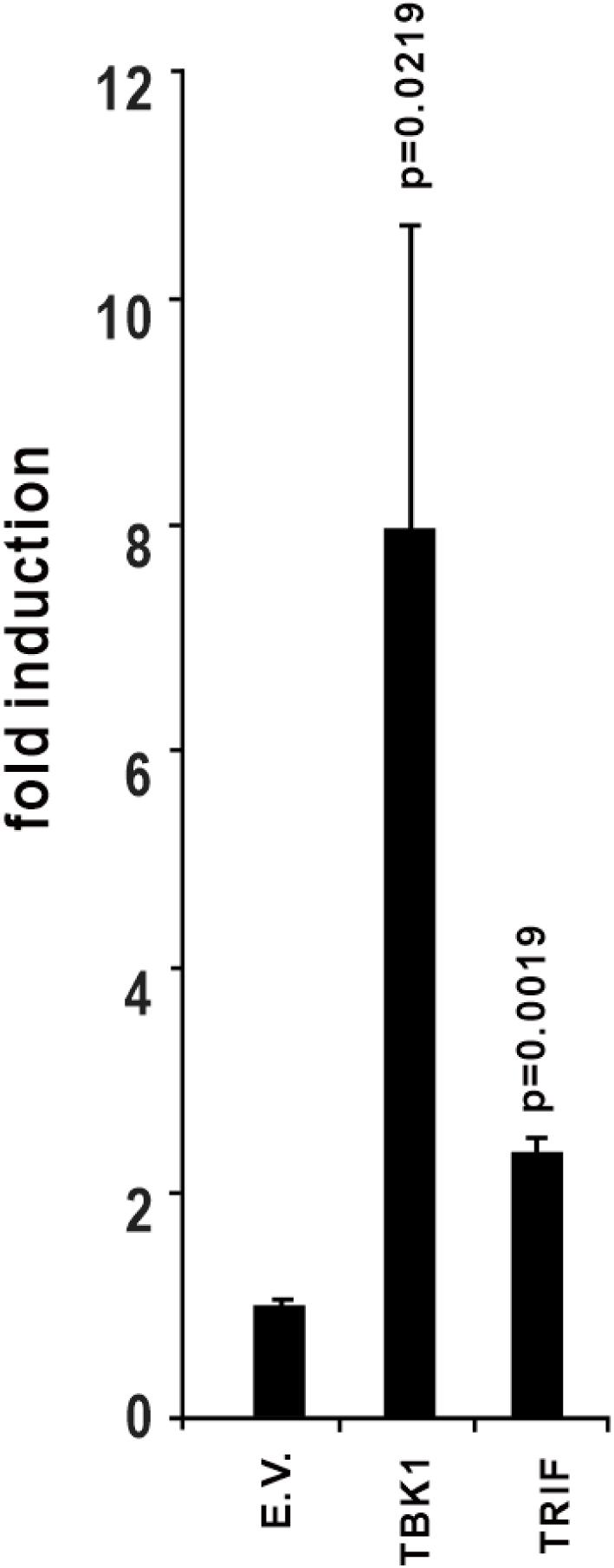
Overexpression of TBK1 or TRIF induces ACE2 expression. TBK1 is an essential activator of RNA-sensing pathway. HEK293 cells were cultured in 24-well plates. 24 hrs later, plasmids as shown above were transfected into HEK293 cells. 36 hrs after transfection, total RNA was isolation for subsequent RT-qPCR analysis. Results in each panel are representative of three independent experiments.

### Virus Infection Induces ACE2 Upregulation

We retried microarray database submitted in NCBI and analysis ACE2 expression levels during virus infection. In lungs from C57Bl/6 mouse infected with 10^2^ PFU SARS-CoV, ACE2 mRNA is significantly increased comparing with the mock group (Figure 4) (Gralinski et al., 2018; Totura et al., 2015). In primary human airway epithelial cells, MERS-CoV infection can obviously increase ACE mRNA levels at 24, 36 and 48 hours post infection (Figure 5). These results clearly reveal that ACE can be upregulated by infection of SARS-CoV and MERS-CoV. Next, we evaluated ACE expression after other RNA virus infection. Data from microarray shows that the ACE2 mRNA level can also be strongly induced in human airway epithelial cells from 4 donors responses to rhinovirus (Figure 5) (Proud et al., 2012). In wild-type H1N1 influenza-infected human bronchial epithelial cells, ACE2 can dramatically increase to about 5-folds higher than the control group (Figure 7) (Shapira et al., 2009). These pieces of evidence indicate that ACE2 is a virus infection responsive gene and its expression can be stimulated by various viruses. Overall, ACE2 expression can be upregulated by virus infection.

**Figure 4.**
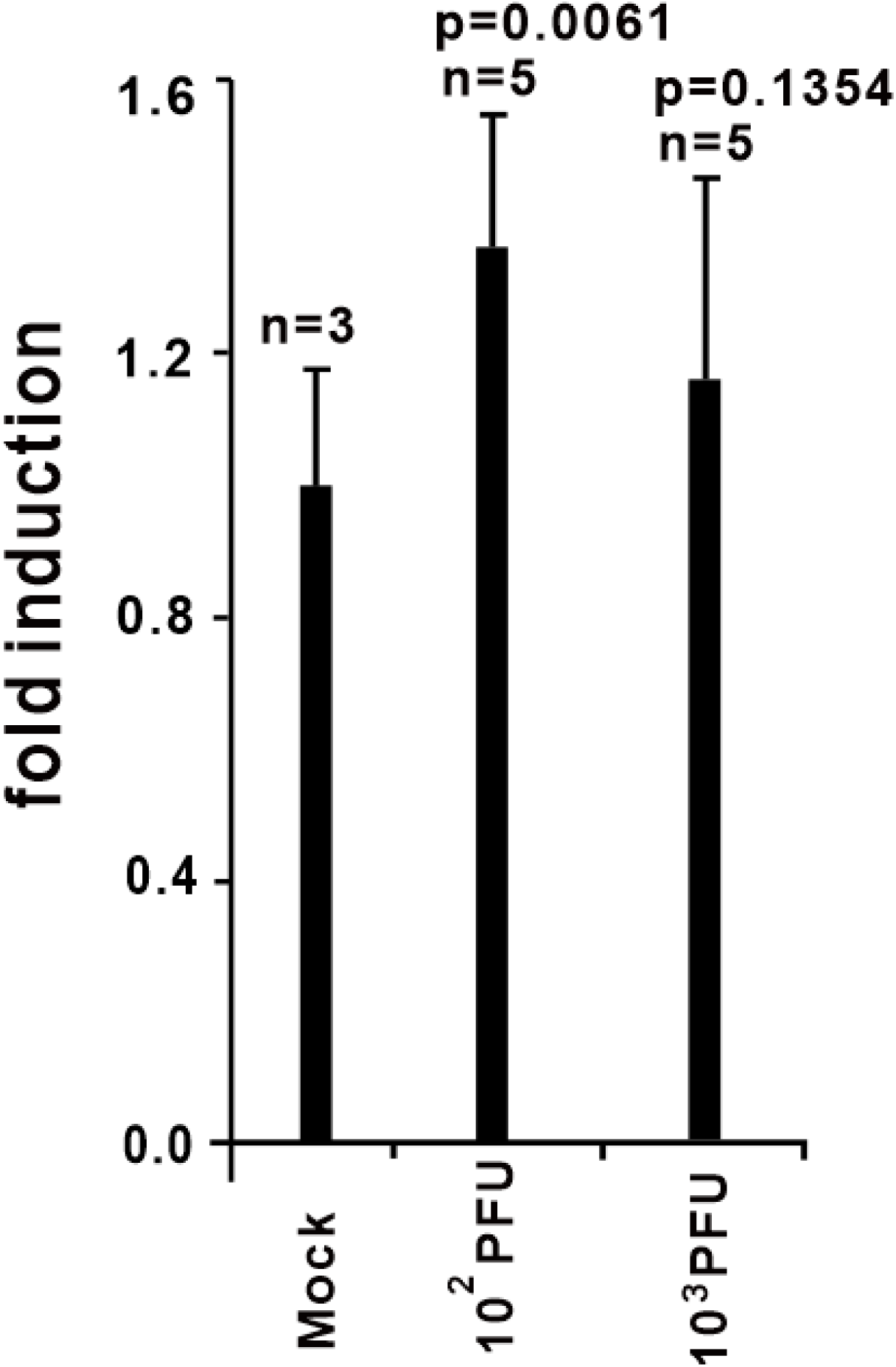
ACE2 expression is upregulated in the lungs from SARS-CoV MA15 infected C57Bl/6 mouse. Intranasal instillation of 10^2^, 10^3^ PFU of SARS-CoV MA15 in 50 µl of PBS or mock-infected with PBS alone, lungs were harvested 24 hrs later and total RNA was isolated and subjected to microarray analysis. ACE2 expression data was retrieved from Gene Expression Omnibus (GEO) microarray database (NCBI Accession: ***PRJNA149057***; GEO: ***GSE33266***).

**Figure 5.**
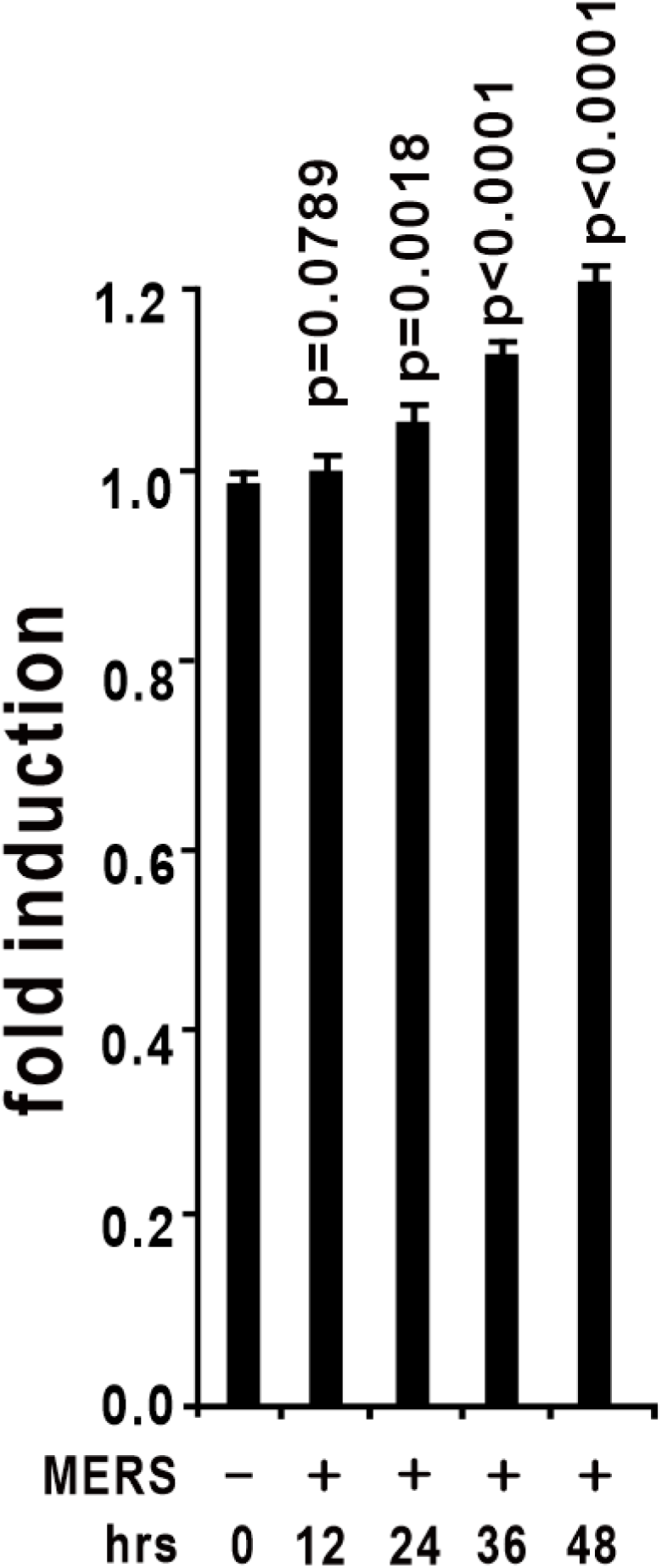
ACE2 expression is upregulated in primary human airway epithelial cells in response to wild type MERS-CoV (icMERS-CoV EMC2012). Human airway epithelial cells were infected with a multiplicity of infection of 5 PFU per cell. Infected samples were collected as indicated above. ACE2 expression data was retrieved from GEO microarray database (NCBI Accession: ***PRJNA391962***; GEO: ***GSE100504***).

**Figure 6.**
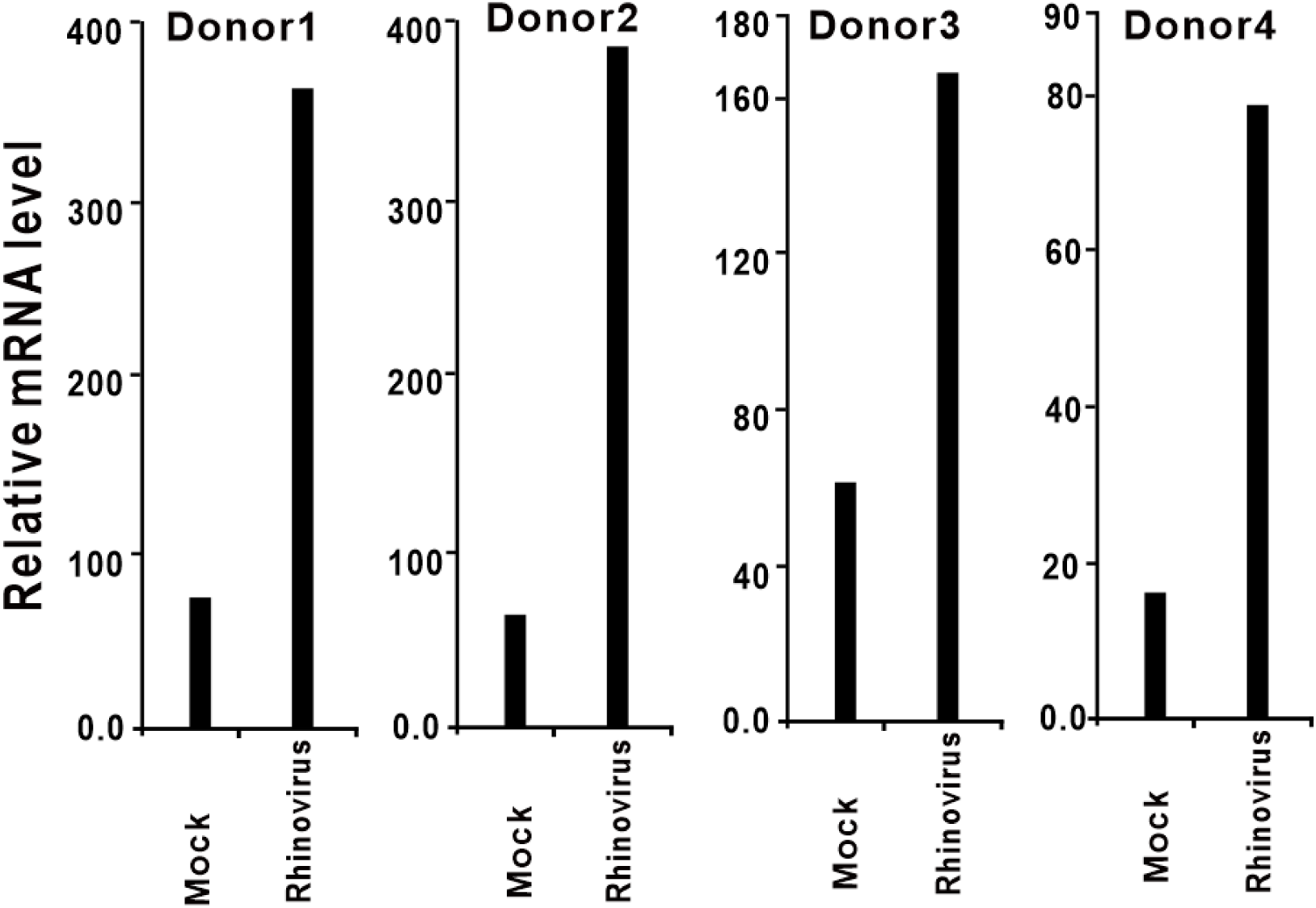
ACE2 expression is upregulated in human airway epithelial cells in response to rhinovirus infection. Primary epithelial cells from 4 donors were exposed to rhinovirus or medium only (used as control). 24 hrs post-infection, gene expression was assessed using Affymetrix U133 plus 2.0 human GeneChips. ACE2 expression data was retrieved from GEO microarray database (NCBI Accession: ***PRJNA137767***; GEO: ***GSE27973***).

**Figure 7.**
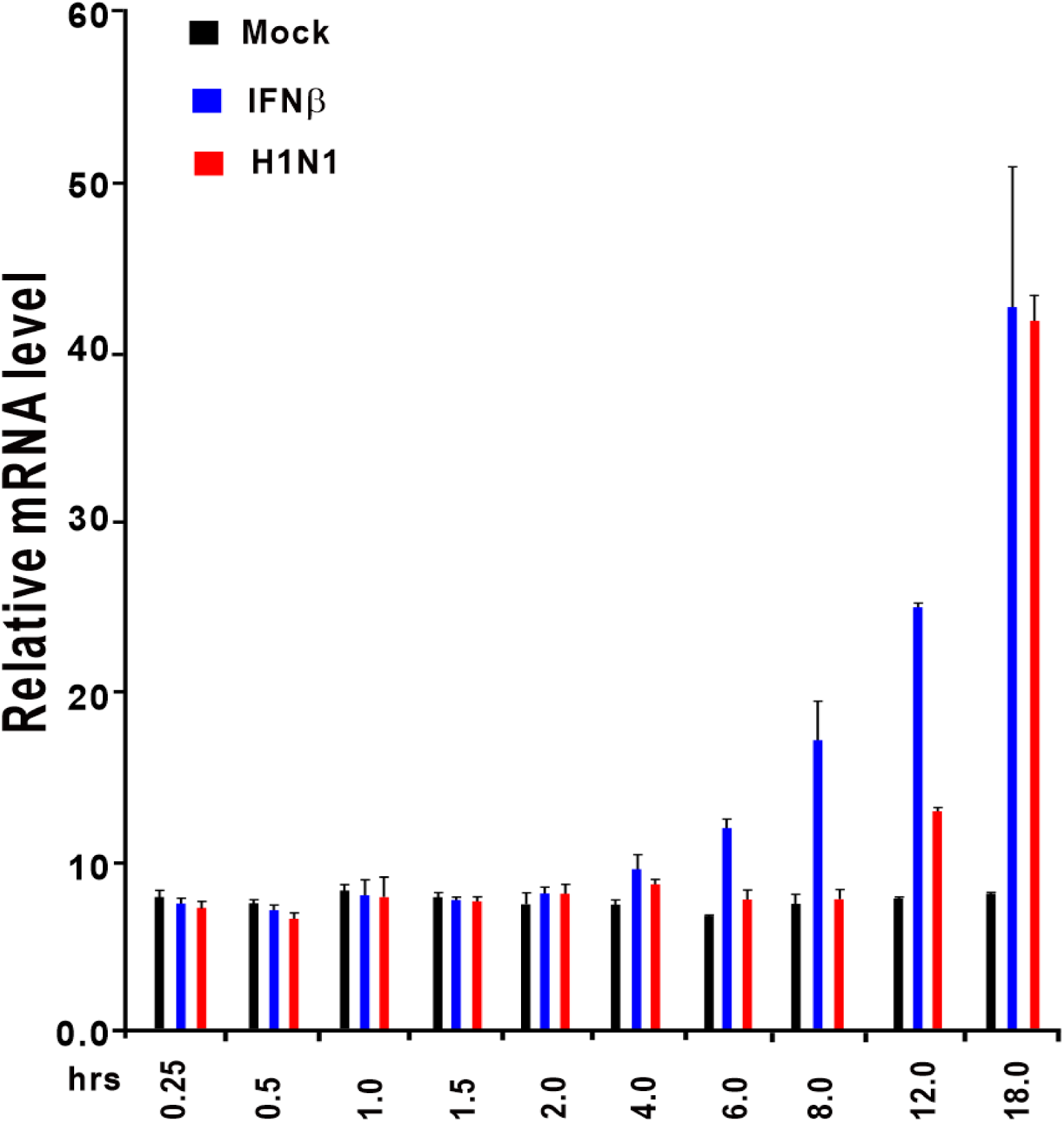
ACE2 expression is upregulated in human bronchial epithelial cells (HBECs) after IFNβ and H1N1 influenza stimulation. HBECs were treated with IFNβ (1000U/ml) or wild type H1N1 influenza (A/PR/8/34). Control samples were incubated with fresh medium under the same conditions. HBECs were harvested at indicated time points (0.25, 0.5, 1, 1.5, 2, 4, 6, 8, 12, and 18 hours post-treatment). Total RNA was extracted and analyzed using the Affymetrix High Throughput Array. ACE2 expression data was retrieved from GEO microarray database (NCBI Accession: ***PRJNA121751***; GEO: ***GSE19392***).

### Inflammatory Cytokines Stimulate ACE2 Expression

We suppose that ACE2 induction by viruses is caused by virus infection-induced inflammatory cytokines. From the microarray database, we observed that ACE2 expression can also be stimulated by type I IFNs including IFNα and IFNβ and type III IFN such as IFNγ. In human bronchial epithelial cells stimulated by IFNβ (1000U/ml), ACE2 is significantly upregulated at 6, 8, 12, and 18 hours post treatment, especially at 18 hours (Figure 7), ACE2 is increased more than 4-folds comparing with the mock group (Shapira et al., 2009). We also found that ACE2 can be significantly induced by IFNγ treatment in human primary keratinocytes (Figure 8) (Swindell et al., 2012). Taken together, ACE2 mRNA is increased in response to inflammatory cytokine stimulation.

**Figure 8.**
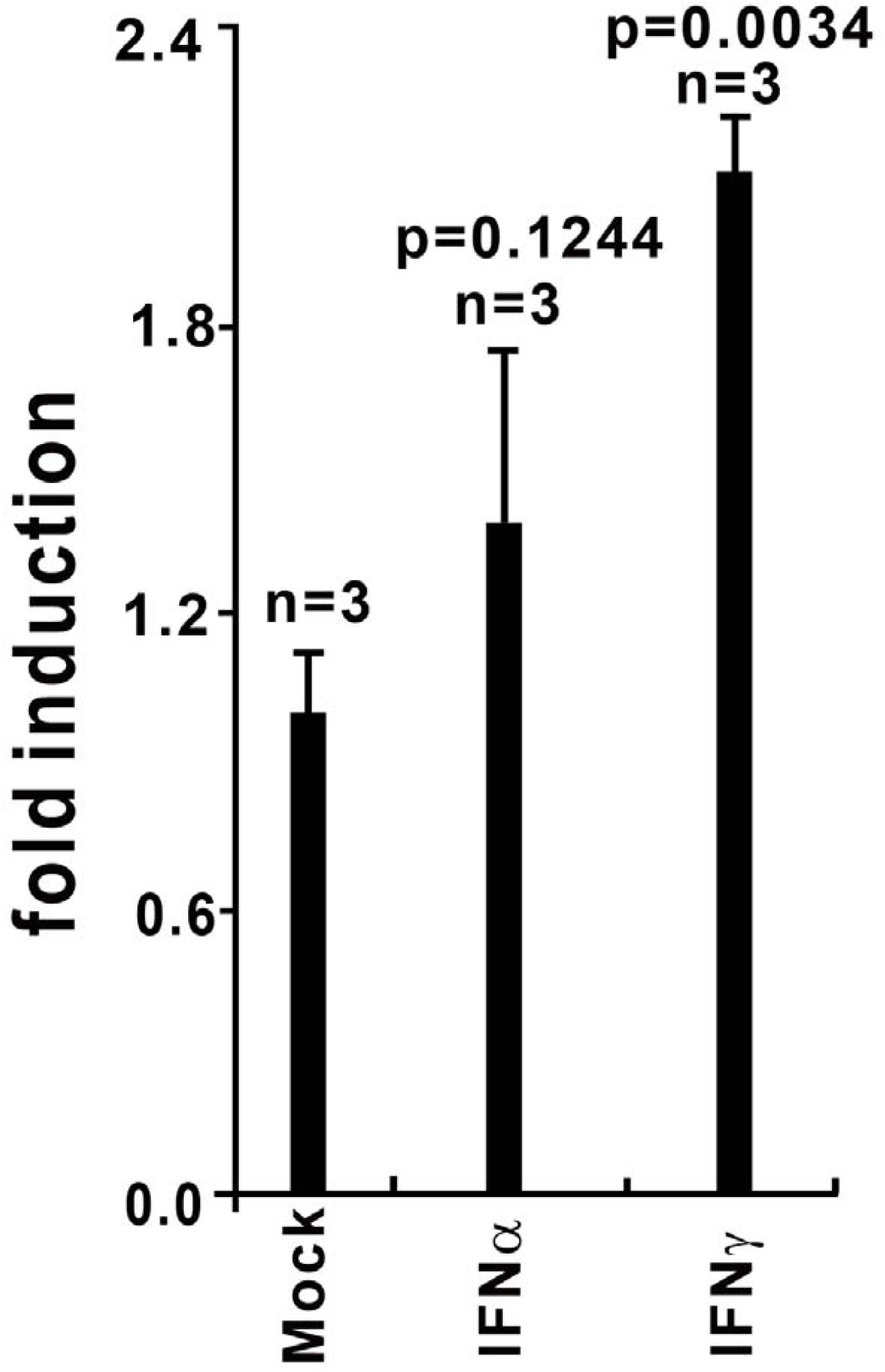
ACE2 expression is upregulated in human primary keratinocytes exposed to IFNα and IFNγ. Primary keratinocytes from 3 donors were either untreated (control) or exposed to cytokines IFNα, IFNγ. 24 hrs after treatment, total RNA was extracted for subsequent microarray analysis. ACE2 expression data was retrieved from GEO microarray database (NCBI Accession: ***PRJNA153023***; GEO: ***GSE36287***).

## Discussion

2019-nCoV infection causes a direful transmit speed and a much unidentified pathological phenomenon. 2019-nCoV utilizes cell membrane receptor for entering into cells and causes infection diseases. Although it is believed that lung is the target organ of 2019-nCoV, lots of non-respiratory symptoms have subsequently been reported, suggesting the involvements of other organs during the disease (Wei et al. 2020). These organs and tissues commonly have high ACE2 expression level such as kidney and small intestine. SARS-nCoV infects multiple cell types in several organs; immune cells and pulmonary epithelium are also the main sites of injury (Gu et al., 2005). Acute kidney injury is a predictor of high mortality in SARS patients (Chu et al., 2005). In the lung, human ACE2 is primarily existed in type II and type I alveolar epithelial cells. ACE2 is mainly expressed in AT2 cells and also detected in AT1 cells, airway epithelial cells, fibroblasts, endothelial cells, and macrophages (Wei et al. 2020).

As the cellular receptor of SARS-CoV and 2019-CoV, ACE2 plays an essential role during these virus infections (Li et al., 2003; Zhou et al., 2020). The ACE2 level is the decisive factor for 2019-nCoV infection (Zhou et al., 2020). These observations drive us to explore whether ACE2 is a virus infection responsive gene. Here we analyzed the database of virus infected- or inflammatory cytokine-stimulated cells or tissues and clearly show that ACE2 can be upregulated by virus infection and inflammatory cytokine-stimulation. However, whether the upregulation by ACE2 by the virus is achieved by virus infection-induced inflammatory cytokinesis is currently unknown. We already notice that the promoter region of ACE2 exists several immune- and cytokine-responsive transcript factor binding sites such as JUN. Next, we will evaluate whether ACE2 is upregulated by JNK signaling cascades. Importantly, the inducibility of ACE2 by inflammatory cytokines also implies that “cytokine storm” caused by 2019-nCoV not only damage host tissues but also may accelerate virus spread (Chen et al., 2020; Huang et al., 2020).

Overall, this study provides the first evidence that SARS-CoV, MERS-CoV and maybe 2019-nCoV can upregulate its cellular receptor ACE2 to facilitate their infection and spread. Thus, reducing ACE2 expression by siRNA or other drugs may be a direction for drug development. Because this cluster of viruses uses the same cellular receptor ACE for entering, thus preparation drugs which can reduce ACE2 expression may be useful for the future emerging infectious diseases. Our study also suggests that people infected by other viruses such as influenza or with high inflammatory cytokines may more susceptible to 2019-nCoV. This study also indicated that targeting the expression of ACE2 is a direction for drug development and infection control.

